# COMMD1 and PtdIns(4,5)P2 interaction maintain ATP7B copper transporter trafficking fidelity in HepG2 cells

**DOI:** 10.1101/572578

**Authors:** Davis J. Stewart, Kristopher K. Short, Breanna N. Maniaci, Jason L. Burkhead

## Abstract

Copper-responsive intracellular ATP7B trafficking is critical to maintain copper balance in mammalian hepatocytes and thus organismal copper levels. The COMMD1 protein binds both the ATP7B copper transporter and phosphatidylinositol (4,5)-*bis*phosphate (PtdIns(4,5)P_2_), while COMMD1 loss causes hepatocyte copper accumulation. Although it is clear that COMMD1 is included in endocytic trafficking complexes, a direct function for COMMD1 in ATP7B trafficking has not been defined. In this study, experiments using quantitative reveal that COMMD1 modulates the copper-responsive ATP7B trafficking through recruitment to PtdIns(4,5)P_2_. Decreased COMMD1 abundance results in loss of ATP7B from lysosomes and the *trans*-Golgi network (TGN) in high copper conditions, while excess expression of COMMD1 also disrupts ATP7B trafficking and TGN structure. Overexpression of COMMD1 mutated to inhibit PtdIns(4,5)P_2_ binding has little impact on ATP7B trafficking. A mechanistic PtdIns(4,5)P_2_-mediated function for COMMD1 is proposed that is consistent with decreased cellular copper export due to disruption of the ATP7B trafficking itinerary and accumulation in the early endosome when COMMD1 is depleted. PtdIns(4,5)P_2_ interaction with COMMD1 as well as COMMD1 abundance may both be important in maintenance of specific membrane protein trafficking pathways.

**SUMMARY:** Quantitative analysis of 3D protein colocalization defines the cellular function of COMMD1 in maintenance of ATP7B copper transporter trafficking fidelity and the importance of PtdIns(4,5)P_2_ in this action.

## INTRODUCTION

Transmembrane and membrane-associated proteins are critical components of the cell plasma membrane as well as organelles, where transport, catalytic activity, signaling and intracellular trafficking must be regulated based on cell physiologic state. Therefore, the proper function and localization of these proteins is critical to the wellbeing of the cell and to the overall health of the organism. The presence and abundance of proteins at the cell surface is regulated by (1) the secretion of proteins to the PM (a process known as exocytosis or anterograde trafficking) and (2) the internalization of proteins at the PM for degradation or recycling (a process known as endocytosis or retrograde trafficking). Proteins following the exocytic pathway leave the *trans*-Golgi network (TGN) and travel directly to the PM in secretory vesicles. Alternatively, endocytic vesicles containing internalized PM proteins quickly target the early endosome (EE). From here, sorting machinery determines the subsequent fate of each cargo protein: targeting for recycling back to the PM, degradation via lysosome, or retrograde routing to the TGN. This sorting machinery involves the coordinated action of numerous regulatory molecules.

The Copper Metabolism Murr1 Domain 1 (COMMD1) protein and COMMD family have gained recent interest for roles in endosomal sorting. COMMD1 is the founding member of the 10 protein COMMD family and was first identified in Bedlington Terriers suffering from canine copper toxicosis. These animals carried a deletion in both alleles of exon 2 of the *Murr1* gene, resulting in a disruption of biliary copper excretion along with chronic hepatitis and cirrhosis similar to Wilson disease in humans (Klomp et al., 2003; Rothuizen et al., 1999; Van De Sluis et al., 2002). COMMD1 quickly became of interest as a regulator of cellular copper levels. Burstein *et al.* (2004) first established a link between COMMD1 and intracellular copper accumulation *in vitro* using mRNA interference in the human embryonic kidney epithelial cell line (HEK293) (Burstein et al., 2004). Additionally, it was shown that siRNA-mediated knockdown of COMMD1 in canine hepatic cells resulted in increased copper retention and sensitivity to excess extracellular copper (Spee et al., 2007).

COMMD1 binds with high specificity to the signaling lipid phosphatidylinositol 4,5-bisphosphate (PtdIns(4,5)_2_) via the conserved C-terminal domain and is stabilized by the formation of higher order oligomers (Burkhead et al., 2009). Furthermore, COMMD1 has also been identified as a factor in endosomal sorting of several other membrane proteins. It was recently reported that COMMD1 formed a novel complex with CCDC22 as well as the previously uncharacterized CCDC93 and C16orf2 (Phillips-Krawczak et al., 2015). This COMMD/CCDC22/CCDC93 (CCC) complex is recruited by FAM21 to endosomes. FAM21 functions in coordination with the Wiskott-Aldrich syndrome protein and SCAR homologue (WASH) complex, retromer, and sorting nexins to facilitate tubulation of early endosomes and receptor trafficking. The authors found that the CCC complex was necessary for ATP7A retrograde trafficking including ATP7A routing to both the PM and to the TGN, indicating other factors may be important to determine the correct pathway (Phillips-Krawczak et al., 2015). The CCC complex in coordination with the WASH complex has also been reported as essential in regulating levels of circulating low-density lipoprotein (LDL) by mediating the endosomal sorting of its receptor, LDLR. Mutations that affect the formation of the CCC complex resulted in mislocalization of LDLR and reduced LDL uptake (Bartuzi et al., 2016).

In humans, copper toxicosis due to Wilson disease is attributed to a loss of function mutation in the copper-transporting ATPase ATP7B (Bull et al., 1993). ATP7B has a dual role in maintaining copper homeostasis: in the liver, ATP7B functions in the TGN where it transports copper into the lumen for biosynthesis of secreted cuproproteins. When cellular copper levels are elevated, ATP7B traffics in vesicles towards the apical membrane to move excess copper into bile for excretion from the body (La Fontaine and Mercer, 2007; Lutsenko et al., 2007). COMMD1 interacts with ATP7B *in vivo* and *in vitro* (Tao et al., 2003), supporting a role in ATP7B regulation. Additionally, siRNA-mediated knockdown of COMMD1 resulted in decreased ATP7B at the TGN in the mouse hepatoma cell line, Hepa 1-6, when cells were treated with a high copper for 24 hours, then exposed to a copper chelator, implying a function in retrograde trafficking (Miyayama et al., 2010). However, the specific function of COMMD1 in ATP7B trafficking, and thus in regulating cellular copper export, remains unclear.

In this study, we tested the hypothesis that the role of COMMD1 in copper homeostasis is through modulation of the ATP7B trafficking itinerary. Quantitative immunofluorescence microscopy was used to monitor ATP7B subcellular localization in the HepG2 hepatoma cell line. COMMD1 impacts ATP7B trafficking in cells, where reduced levels of COMMD1 result in a loss of ATP7B colocalized with lysosomes as well as at the TGN, and an accumulation of ATP7B associated with retromer at the EE. Overexpression of COMMD1 also promoted mis-localization of ATP7B, while expression of COMMD1 variants defective in PtdIns(4,5)P_2_ binding diminished this effect. Taken together, these data illustrate the functional role of COMMD1 to maintain ATP7B trafficking fidelity and indicate the importance of COMMD1 interaction with PtdIns(4,5)P_2_ to exert effects on ATP7B. These observations reveal new potential functional roles for COMMD proteins and specific interactions with signaling lipids in fidelity of membrane protein trafficking.

## RESULTS

### Quantitative colocalization monitors ATP7B redistribution in response to copper exposure

A quantitative colocalization approach was used to monitor the copper-responsive trafficking itinerary for ATP7B in HepG2 cells. ATP7B is known to localize at the TGN where it functions to transport copper into the lumen for biosynthesis of secreted cuproproteins (Lutsenko et al., 2008). When copper levels increase, ATP7B traffics in vesicles to lysosomes and toward the apical membrane to facilitate copper sequestration or efflux (Polishchuk et al., 2014), though it is expected that some must remain at the TGN for continued cuproprotein synthesis. Polishchuk and co-workers established a lysosomal location for non-TGN ATP7B under high Cu conditions (Polishchuk et al., 2014). Therefore, we analyzed ATP7B colocalization with lysosomes and the TGN in HepG2 cells under varying copper treatments. Pearson and Manders (M1 and M2) coefficients were calculated to define ATP7B colocalization in individual cells. As previously described, Pearson values are a useful measure of probe association where as Manders is a direct measure of the fraction of colocalized probe fluorescence (Dunn et al., 2011). Correlation coefficients for each pair of proteins were plotted and curves were fitted to visualize the distribution of values in each treatment and identify changes in ATP7B colocalization with specific compartment markers in response to Cu levels.

Representative images with approximately median colocalization values are shown in **(Fig. 1A**,**B)**. Both Manders values indicate that the ratio of ATP7B localized with the TGN remained constant under all copper treatments **(Fig. 1C**,**D)**. The Pearson coefficients also indicate the variance in relation to the TGN does not change with the exception of a slight insignificant increase under low copper **(Fig. 1E)**. In contrast, the ATP7B lysosomal pool showed a significant response to Cu, with increased mean values for both Manders and Pearson coefficients when treated with 10, 100, and 200 µM copper and decreased values when treated with the copper chelator tetrathiomolybdate (TTM) compared to basal media **(Fig. 1F-H)**. This, along with the observation that localization remained constant at the TGN indicates that ATP7B synthesis is likely upregulated under high copper, and that the increase in the pool is shifted toward the lysosomes. Under low copper conditions excess ATP7B leaves lysosomes.

**Figure 1.**
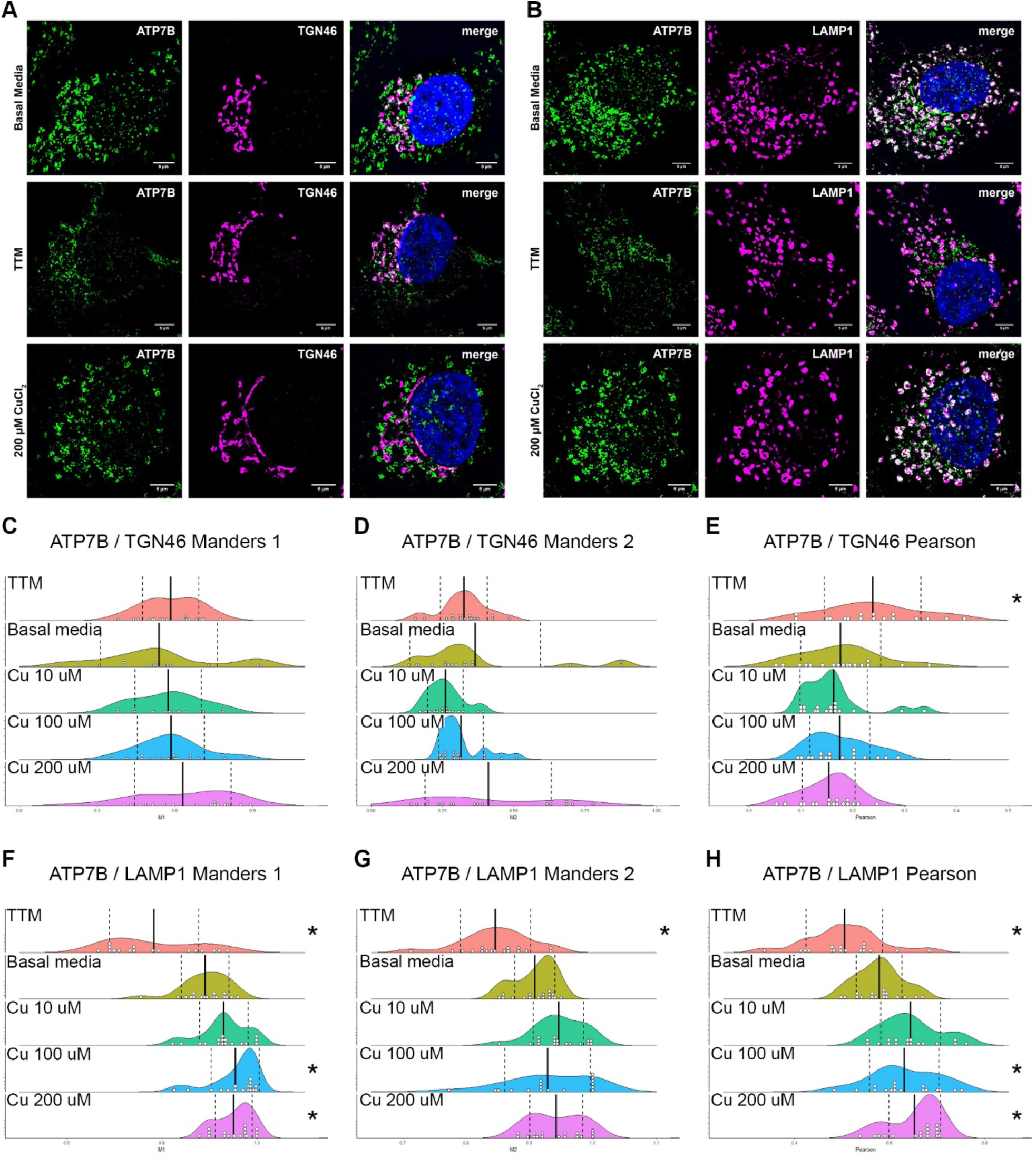
Colocalization of ATP7B with TGN46 or LAMP1. Cells were left untreated (basal media) or treated with low copper (10 μM TTM), 10, 100, or 200 μM copper. Merged image shows ATP7B in green, TGN46 **(A)** or LAMP1 **(B)** in magenta and the nucleus in blue; pixel overlap is shown in white. 3D colocalization analysis produced Manders and Pearson correlation coefficients for ATP7B and TGN46 **(C-E)** or LAMP1 **(F-H)**. Graphs show density plots with treatment means (solid line) and standard deviation (SD, dashed lines) shown. Individual cell values are indicated as a white circle (binned at 0.005). Treatment means were tested against the untreated cells (basal media) using Dunnett’s method. “*” indicates a p-value < 0.05 between the control and the treatment means.

Taken together, these data demonstrate that copper-responsive ATP7B distribution can be quantified by 3-D colocalization analysis. Furthermore, ATP7B localization is not binary—the data indicate continuous trafficking between various cellular compartments. These results support a trafficking itinerary where the cell responds to changes in copper levels by adjusting the proportion of ATP7B localized at a specific target compartment: when cellular copper levels decrease, the lysosomal pool of ATP7B decreases; when copper levels increase through addition of 10, 100, or 200 µM Cu to the media, ATP7B is directed to the lysosome.

### COMMD1 is a modifier of ATP7B localization

Bedlington Terriers with a loss-of-function mutation in both alleles of COMMD1 suffer copper toxicosis due to an impaired ability to excrete biliary copper (Van De Sluis et al., 2002). Furthermore, liver-specific COMMD1 knockout mice fed a copper-enriched diet had increased intrahepatic copper levels compared to control mice (Vonk et al., 2011). Therefore, we determined the impact of COMMD1 depletion by siRNA on copper-responsive ATP7B trafficking.

We found that ATP7B colocalization with the TGN increased under low copper conditions in cells with decreased COMMD1 expression as compared to control cells **(Fig. 2A-C)**. However, M1 remained unchanged, indicating that the proportion of total ATP7B localized with the TGN increased, while the amount of ATP7B at the TGN did not. When treated with 200 µM copper, both the amount of ATP7B (M1) and the proportion of ATP7B (M2) localized at the TGN decreased significantly in cells with reduced COMMD1 expression. The Pearson correlation coefficient for ATP7B with lysosomes decreased in cells with depleted COMMD1 under low copper conditions while M1 and M2 both remained unchanged **(Fig. 2D-E)**. Pearson, M1, and M2 for ATP7B with lysosomes all showed a significant decrease in 200 µM copper-treated cells with reduced COMMD1. This indicates that under high copper conditions, both the amount of ATP7B at the lysosomes and the proportion of the ATP7B pool that is associated with lysosomes decreases when COMMD1 is less abundant.

**Figure 2.**
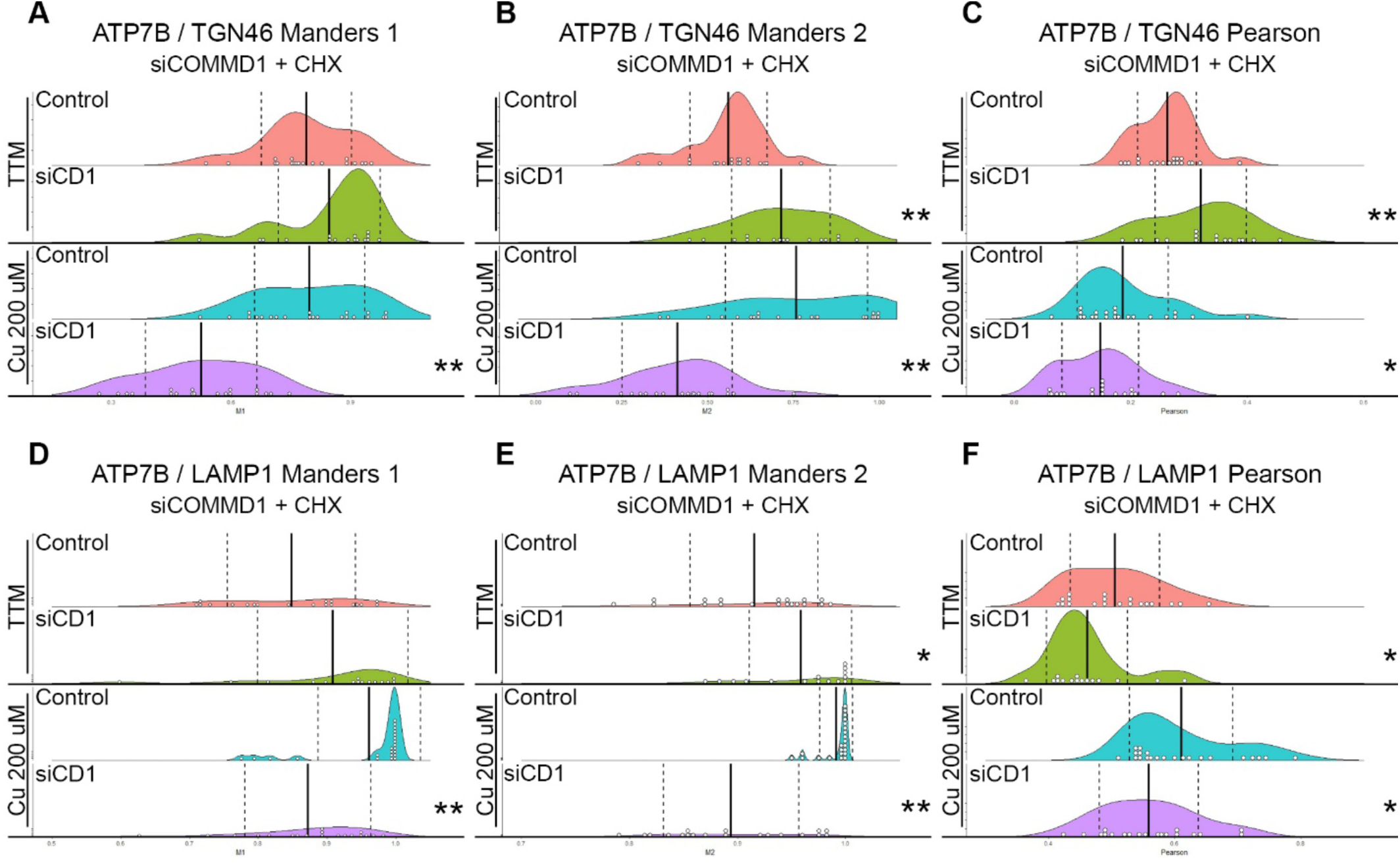
Colocalization of ATP7B with with TGN46 or LAMP1 in cells with reduced COMMD1 expression. Cells were transfected with COMMD1 (siCD1) or nontarget (control) siRNA and treated with TTM (low Cu) or 200 μM Cu and Cycloheximide for the last hour. 3D colocalization analysis produced a Manders **(A, B, D**, and **E)** and Pearson **(C** and **F)** correlation coefficient for each cell. Graphs show density plots with treatment means (solid line) and SD (dashed line). Individual cell values are indicated as a white circle (binned at 0.005). A t-test was used to compare COMMD1 knockdown to the control for each copper treatment. “*” indicates a p-value < 0.05 “**” indicates a p-value < 0.005.

Replicate wells were additionally treated with chloroquine and MG132 to prevent protein degradation. Chloroquine functions to prevent endosomal acidification thereby preventing fusion of endosomes and lysosomes. Furthermore, it increases lysosomal pH, inhibiting protein degradation. MG132 was added to prevent degradation via proteasomes. Under these conditions, colocalization between ATP7B and lysosomes was not significantly different under high or low copper conditions in cells with depleted COMMD1 as compared to the control. ATP7B colocalization with the TGN was also not significantly different in cells treated with high copper. However, when treated with TTM, cells with depleted COMMD1 showed a significant (P=0.0195) increase in the Pearson coefficient, indicating increased association between ATP7B and the TGN **(Fig. 3)**. However, Manders 1 and 2 were not significantly different and therefore the proportion of ATP7B localized with TGN46 and vice-versa did not change.

**Figure 3.**
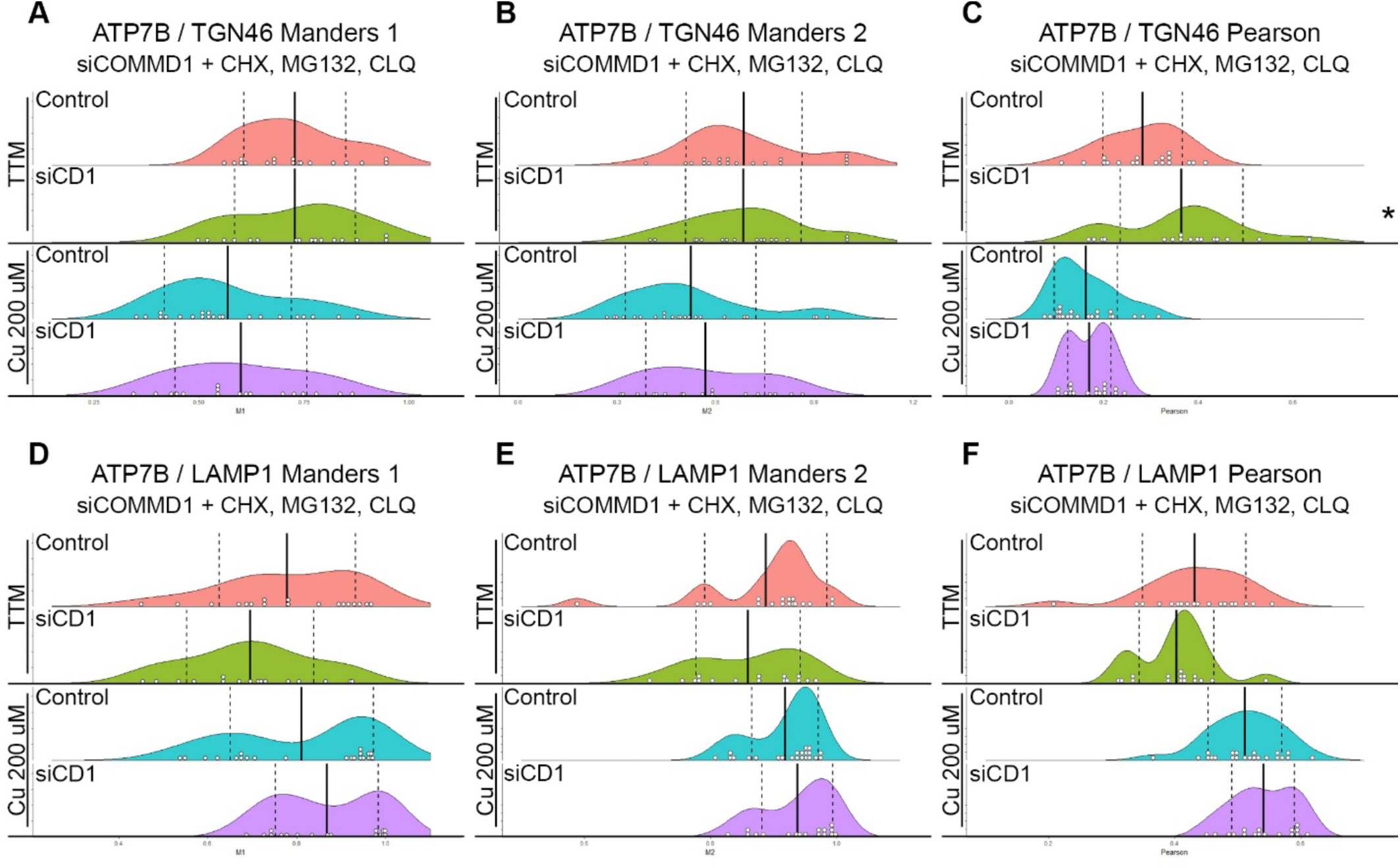
Colocalization of ATP7B with with TGN46 or LAMP1 in cells with reduced COMMD1 expression. Cells were transfected with COMMD1 (siCD1) or nontarget (control) siRNA and treated with TTM (low Cu) or 200 μM Cu and Cycloheximide with the addition of MG132 and Chloroquine (CLQ) for the last hour. Manders **(A, B, D**, and **E)** and Pearson **(C** and **F)** correlation coefficients were generated for each cell and the graph shows density plots with treatment means (solid line) and SD (dashed line). Individual cell values are indicated as a white circle (binned at 0.005). A t-test was used to compare COMMD1 knockdown to the control for each copper treatment. “*” indicates a p-value < 0.05.

Taken together, these results indicate that COMMD1 has an important role in maintaining the ATP7B trafficking itinerary. In cells with reduced COMMD1 expression and treated with TTM, the amount of ATP7B at the TGN remained constant while the proportion of the cellular pool increased, indicating a decrease in the trafficking fraction of ATP7B. The lysosomal pool remained unchanged under these same conditions. Alternatively, when COMMD1-depleted cells were treated with high copper, the amount of ATP7B associated with the both the TGN and lysosome significantly decreased as did the proportion of cellular ATP7B associated with these compartments. This indicates that ATP7B trafficking induced by excess cellular copper levels is altered by depletion of COMMD1, with ATP7B potentially trapped in another compartment. When fusion of endosomes and lysosomes is blocked with chloroquine, the observed shifts in ATP7B distribution are lost. This suggests that COMMD1 is important to direct ATP7B to the recycling pathway in the EE. Thus, when COMMD1 is knocked down, ATP7B may be directed to the lysosomal degradation pathway, or it might be trapped in the endosome.

### ATP7B accumulates in endosomes in COMMD1-depleted cells

The Burstein group reported that COMMD1 forms a novel complex with coiled-coil domain-containing protein 22 (CCDC22), coiled-coil domain-containing protein 93 (CCDC93) and C16orf62. They suggested this COMMD/CCDC22/CCDC93 (CCC) complex is recruited by FAM21 to endosomes (Phillips-Krawczak et al., 2015). FAM21 functions in coordination with the Wiskott-Aldrich syndrome protein and SCAR homologue (WASH) complex, retromer, and sorting nexins to facilitate tubulation of early endosomes and receptor trafficking. Thus, we examined ATP7B localization with VPS35, the core component of retromer, in response to attenuated COMMD1. HepG2 cells were transfected with siCOMMD1 or control siRNA and treated with TTM or CuCl_2_ as above. COMMD1-depleted cells grown in TTM showed a dramatic increase (p-value < 0.0001) in ATP7B:VPS35 colocalization for all three coefficients **(Fig. 4)**. In cells treated with 200 µM CuCl_2_, COMMD1 attenuation did not significantly increase the Pearson coefficient, however both Manders 1 and 2 were again dramatically increased. ATP7B accumulation with VPS35 in these conditions indicates that COMMD1 is important for ATP7B exit from the early endosome and further implicates COMMD1 in ATP7B retrograde trafficking.

**Figure 4.**
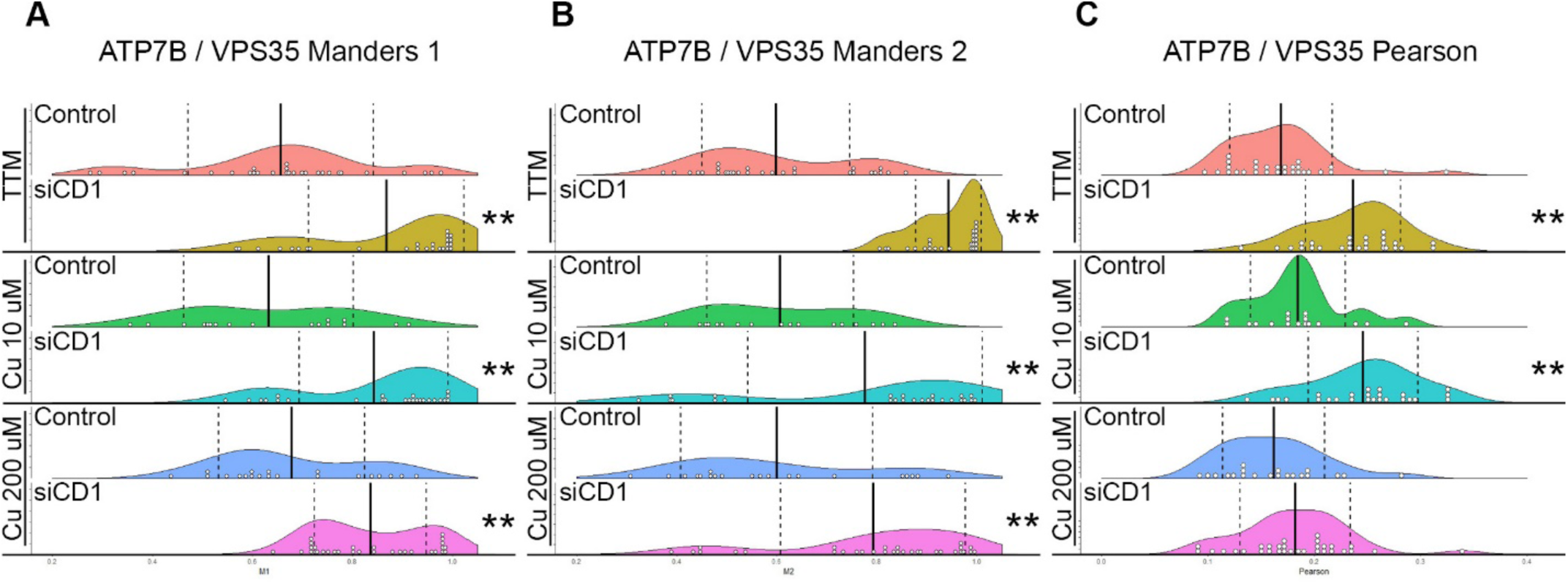
Colocalization of ATP7B with the retromer subunit VPS35 in cells with reduced COMMD1 expression. Cells were transfected with COMMD1 siRNA (siCD1) or nontarget (control) siRNA and treated with TTM (low Cu), 10 μM Cu, or 200 μM Cu and Cycloheximide for the last hour. 3D colocalization analysis produced a Manders **(A** and **B)** and Pearson **(C)** correlation coefficient for each cell. Graphs show density plots with treatment means (solid line) and SD (dashed line). Individual cell values are indicated as a white circle (binned at 0.005). A t-test was used to compare COMMD1 knockdown to the control for each copper treatment. “**” indicates a p-value < 0.005.

### Knockdown of COMMD1 results in an increase in the total pool of ATP7B

Previous studies in multiple cell lines have demonstrated that COMMD1 knockdown by RNA interference results in increased cellular copper levels (Burstein et al., 2004; Spee et al., 2007). However, the interaction of COMMD1 attenuation with ATP7B abundance has not been documented. Using siRNA as in trafficking experiments, we knocked COMMD1 down to varying degrees then assessed the amount of ATP7B present. We found that as COMMD1 levels were reduced, the cellular pool of ATP7B increased **(Fig. 5)**. This inverse correlation fit a linear model with an R^2^ value of 0.87. This data suggests that decreased COMMD1 does not necessarily result in loss of ATP7B, despite apparent loss of copper export. These results are consistent with loss of copper export in Wilson Disease patients with mutations that impact the trafficking itinerary of ATP7B. These observations also indicate that COMMD1 is not likely responsible for preventing ATP7B degradation; but it is instead involved in directing ATP7B to the correct recycling pathway.

**Figure 5.**
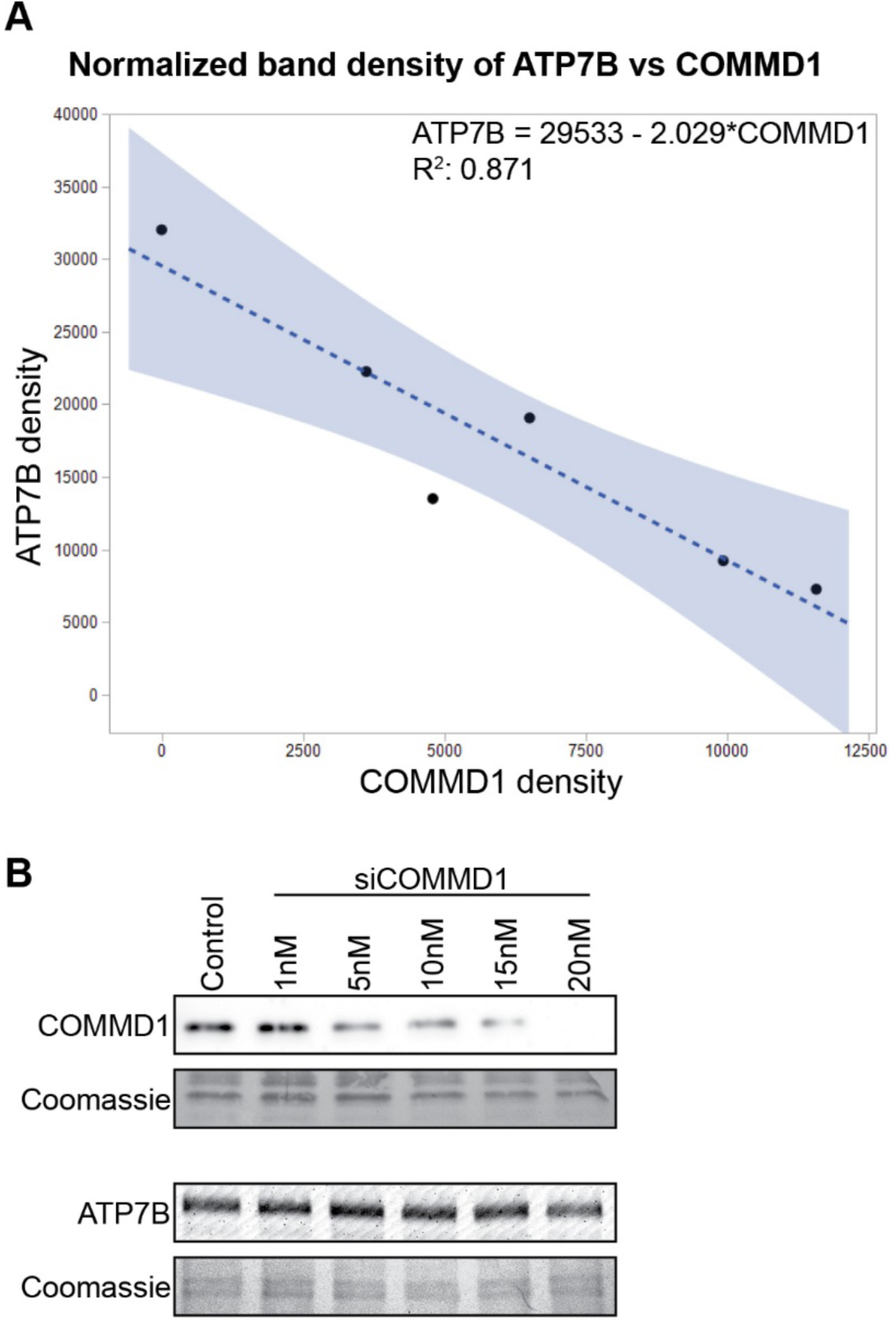
Linear regression of normalized band density of ATP7B vs COMMD1 in cells with reduced COMMD1 **(A)**. HepG2 cells were treated with control siRNA or with 1, 5, 10, 15, or 20 nM siRNA to gradually deplete COMMD1 levels. ATP7B and COMMD1 abundance was assessed by densitometry analysis of western blots with band density normalized to the Coomassie stained blot **(B)**. Statistical analysis shows a negative linear correlation between COMMD1 and ATP7B abundance.

### PtdIns(4,5)P_2_ binding by COMMD1 is important to modulate ATP7B trafficking

Since decreased cellular levels of COMMD1 modulated ATP7B trafficking in a dose-dependent manner, we tested the hypothesis that excess COMMD1 would also impact ATP7B localization. Using an inducible system to increase cellular COMMD1 levels, cells overexpressing wild type COMMD1 under low copper conditions had no significant change in ATP7B colocalization at the TGN. Under high copper conditions, overexpression of wild type COMMD1 resulted in a significant decrease in the Pearson coefficient for ATP7B and the TGN, indicating an impact on trafficking when copper levels are high **(Fig. 6)**.

**Figure 6.**
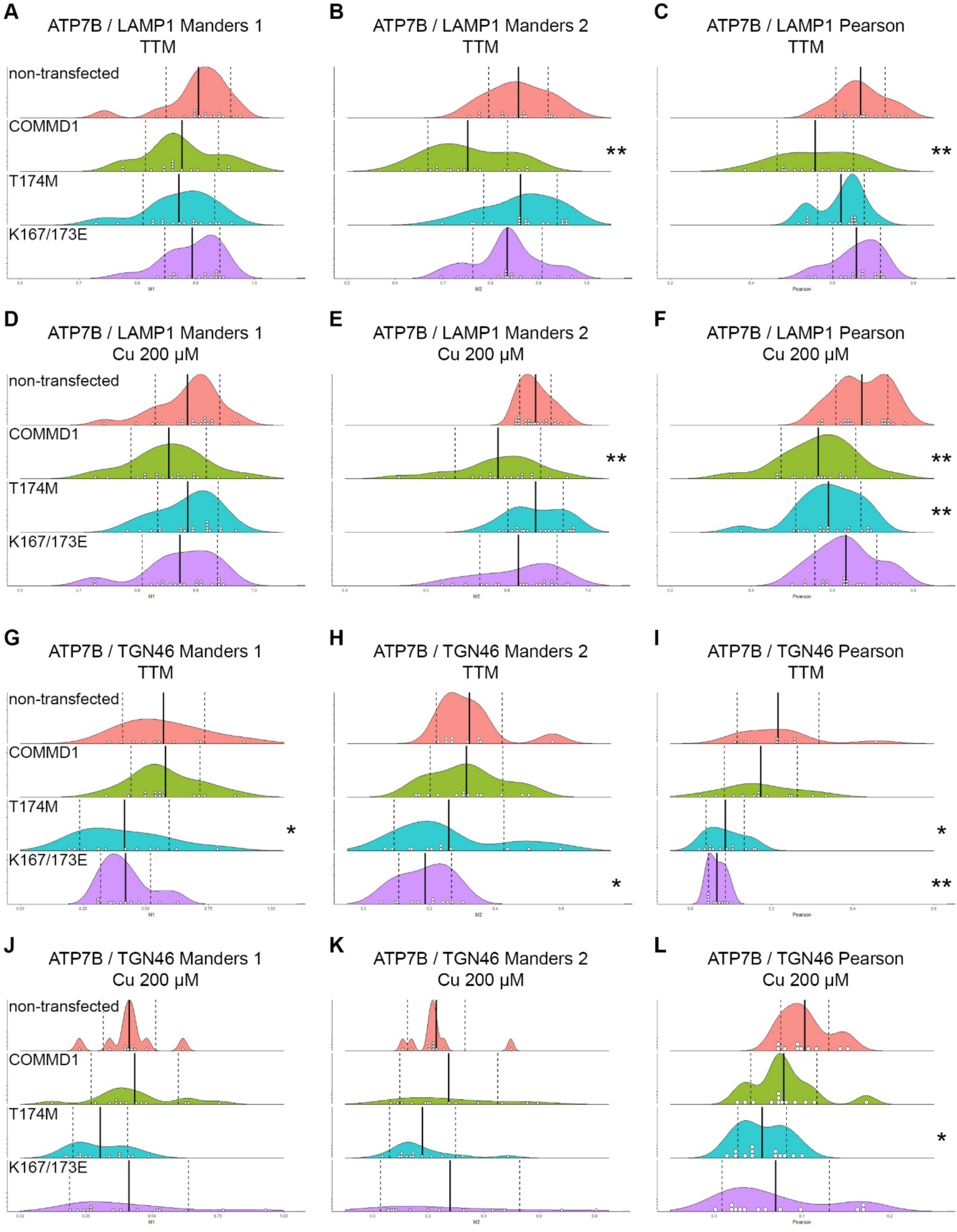
Colocalization of ATP7B with LAMP1 or TGN46 in cells misexpressing COMMD1. Cells were transfected with one of three COMMD1 variants: GFP-COMMD1, GFP-COMMD1 T174M and GFP-COMMD1 K167/173E. Cells were then treated with TTM (low Cu) or 200μM Cu for 9 hours with cycloheximide added for the last hour. 3D colocalization analysis was used to produce a Menders and Pearson correlation coefficient for individual cells. Treatment means (solid line) and SD (dashed line) are shown. Individual cell values are indicated as a white circle (binned at 0.005). “*” indicates a p-value < 0.05 “**” indicates a p-value < 0.005.

A study analyzing at atypical phenotypes of Wilson disease revealed a patient with a missense mutation in COMMD1 resulting in an amino acid change of threonine 174 to methionine (T174M). This patient exhibited unusually severe early onset Wilson disease (Gupta et al., 2010). The T174M mutation is in close proximity to two positively charged residues K167 and K173 **(Fig. 7A)**, which are key targets for PtdIns interaction based on structural models of COMMD1 (Burkhead et al., 2009; Healy et al., 2018). We analyzed COMMD1 wild type and mutant affinities for a soluble PtdIns(4,5)_2_ by intrinsic fluorescence quenching and observed a small decrease in *in vitro* PtdIns(4,5)_2_ affinity for T174M compared with wild type, and a substantial (5-fold) decrease in PtdIns(4,5)_2_ affinity in the K167E/K173E mutant **(Fig. 7B)**. In cells, overexpression of either T174M and K167E/K173E mutants resulted in a significant reduction in the Pearson coefficient for ATP7B with the TGN under TTM or high copper treatments **(Fig. 6I**,**L)**. The Manders M1 and M2 distributions were also shifted down **(Fig. 6G**,**H**,**J**,**K)**; however, only M1 (TGN46_colocalized_/TGN46, or the amount of ATP7B localized at the TGN) decreased significantly. Similar to the wild type, expression of COMMD1 T174M induced a significant decrease in Pearson for ATP7B:lysosomes under high copper but only a minimal shift in the distribution under low copper conditions **(Fig. 6C**,**F)**. However, expression of COMMD1 K167E/K173E had no effect on the localization of ATP7B under either treatment as compared to non-transfected cells **(Fig. 6A**,**B**,**D**,**E)**, indicating that PtdIns(4,5)_2_ binding is important in COMMD1 modulation of ATP7B trafficking.

**Figure 7.**
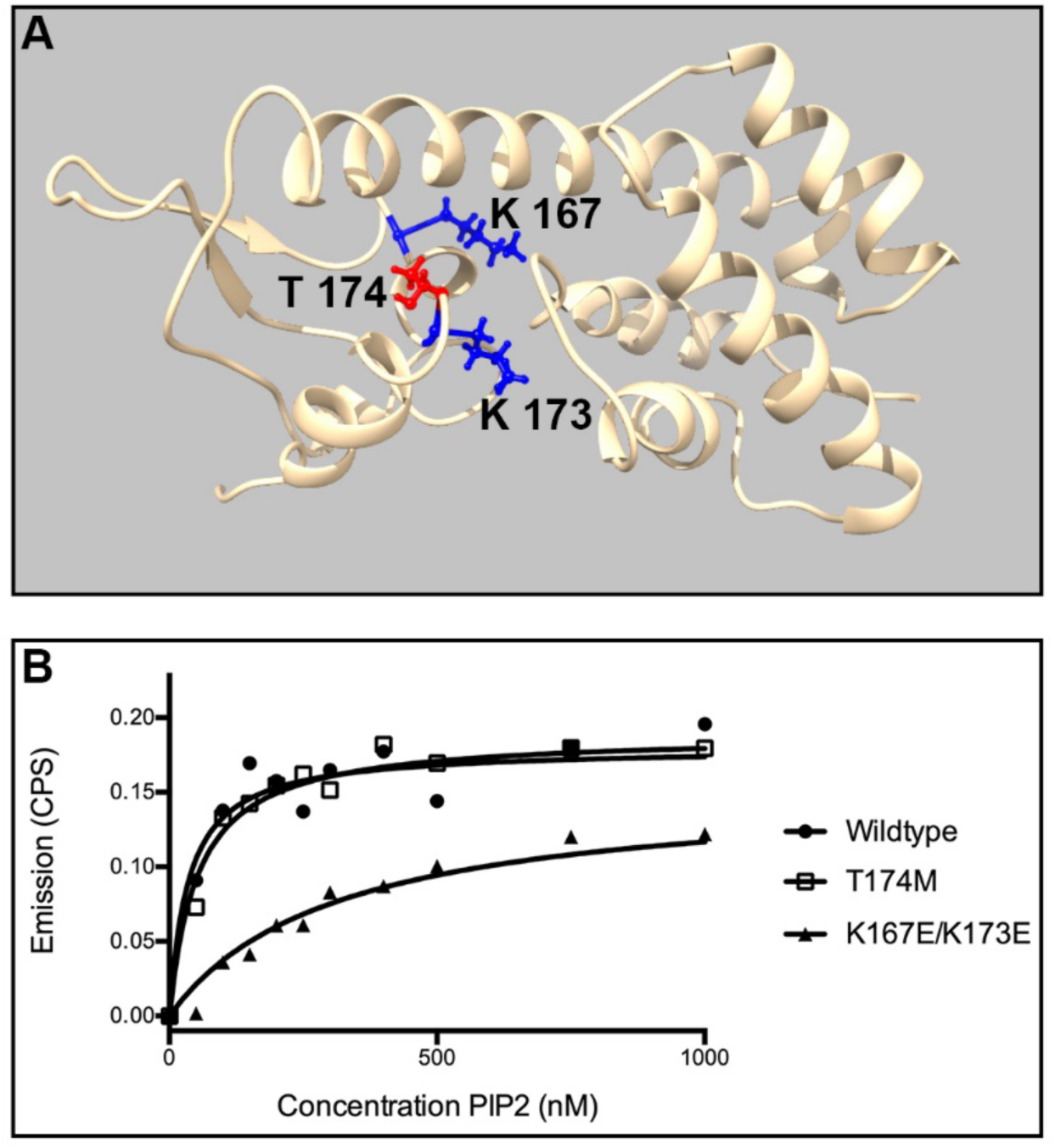
COMMD1-PIP2 Interaction Measured by Fluorescence Quenching of All Aromatics. **(A)** Structural model of COMMD1 showing the close proximity of T174 to the positively charged residues K167 and K173. **(B)** Each line represents the change in fluorescence emission measured at 332nm of one COMMD1 variant with increasing amounts of PIP2 when excited to 280nm. To generate a saturation curve from raw quenching data the formula y=1-(f1/f0) was used, where f1 is the emission in counts per second at a given concentration, and f0 is the emission in counts per second of the first data point where concentration is zero. The Prism (Graphpad) nonlinear regression functions One site Total and One site Specific binding curves were compared for best fit in and it was determined One Site Specific was the best fit for all data sets. One site Specific saturation curves were then generated and yielded K_d_=40.43nM B_max_=0.1871 R^2^=0.9177 for the wildtype, K_d_=62.52 nM B_max_=0.2013 R^2^=0.9788 for T174M and K_d_=284.1 nM B_max_=0.01532 R^2^=0.9721 for K167/173E (n=3) for all data points. The X-axis shows points to 1000 nM for visual display of saturation points, though all data points to 8500 nM were included for calculations.

### Competition for PtdIns(4,5)P_2_ modifies ATP7B trafficking, while global modulation of PtdIns(4,5)P_2_ disrupts critical homeostatic trafficking

COMMD1 binds to PtdIns(4,5)_2_ with high specificity via the conserved C-terminal domain (Burkhead et al., 2009). Therefore, we tested the hypothesis that modification to cellular PtdIns(4,5)_2_ levels will affect ATP7B localization by overexpression of PtdIns(4,5)_2_-modifying enzymes. ADP-ribosylation factor (ARF) 6 regulates trafficking between endosomes and the plasma membrane. Aikawa and Martin, (2003) found that expression of the constitutively active GTPase-defective ARF6^Q67L^ mutant redistributes PtdIns(4,5)_2_ from the plasma membrane to endosomes. If COMMD1 is recruited to endosomes by PtdIns(4,5)_2_, it is expected that more ATP7B would remain in the recycling pool. Alternatively, the overexpression of a PtdIns(4,5)_2_-5-phosphatase should deplete cellular PtdIns(4,5)_2_ levels, resulting in more ATP7B being directed towards degradation. Finally, the Pleckstrin homology domain (PH domain) of Phospholipase C delta has also been shown to bind with high specificity to PtdIns(4,5)_2_ (Várnai and Balla, 1998). Though the binding affinity of the PH domain may be lower than that of COMMD1, PH-domain overexpression might compete with COMMD1 for PtdIns(4,5)_2_ binding sites and shift the distribution of the ATP7B. Thus, to determine how modulation of PtdIns(4,5)_2_ affected ATP7B localization, HepG2 cells were transfected with plasmids to express ARF6^Q67L^, PtdIns(4,5)-5-phosphatase, PLCδ PH domain-GFP fusion, or GFP as a control and treated cells with TTM or Cu as above.

In cells expressing ARF6^Q67L^ under low copper (TTM) conditions, both the Pearson and M1 coefficients ATP7B/TGN46 colocalization were significantly decreased as compared to the control cell population **(Fig. 8 A-C)**. When treated with 200 µM Copper, there was an increase in M2, but M1 and Pearson showed no significant change **(Fig. 8D-F)**. There was no change in colocalization between ATP7B and LAMP1 in cells expressing ARF6^Q67L^ with high or low copper conditions **(Fig. 8G-L)**. Cells expressing PtdIns(4,5)-5-phosphatase lost all TGN46 signal **(Fig. 8A-F)**. This is likely due to an increase in PI(4)P at the *trans-*Golgi. PI(4)P is known to recruit clathrin adaptor AP-1 to the *trans-*Golgi (Wang et al., 2003), which functions to sort proteins for trafficking from the *trans-*Golgi to endosomes and lysosomes. Therefore, it is likely TGN46 anterograde trafficking was dramatically increased, transporting all TGN46 to the endosomes and eventually the lysosome. This likely also impacted protein sorting and trafficking for many other cargo proteins, including ATP7B. These observations suggest that artificially modulating PtdIns(4,5)P_2_ levels in cells is overall deleterious and impacts broad cellular functions. Thus, this data was not used to assess how interactions between COMMD1 and PtdIns(4,5)_2_ affect ATP7B trafficking. Expression of the PLCδ PH domain (fused to GFP) had no significant effect on ATP7B localization with the lysosome and no significant effect on localization with the *trans-* Golgi under high copper conditions **(Fig. 8D-F**,**J-L)**. However, overexpression of PH-GFP resulted in an ATP7B:TGN M2 **(Fig. 8H)** that was significantly higher than the control non-transfected cells, indicating that the distribution of the ATP7B pool had shifted towards the *trans*-Golgi, supporting a role for PtdIns(4,5)P_2_ in ATP7B trafficking. The PH domain was observed at intracellular vesicles and at the plasma membrane in these experiments with HepG2 cells **(Fig. 9)**. This indicates that PtdIns(4,5)_2_ may have a regulatory role at locations other than the plasma membrane, consistent with the observation that COMMD1-PtdIns(4,5)P_2_ interaction is important for COMMD1 to modulate ATP7B location.

**Figure 8.**
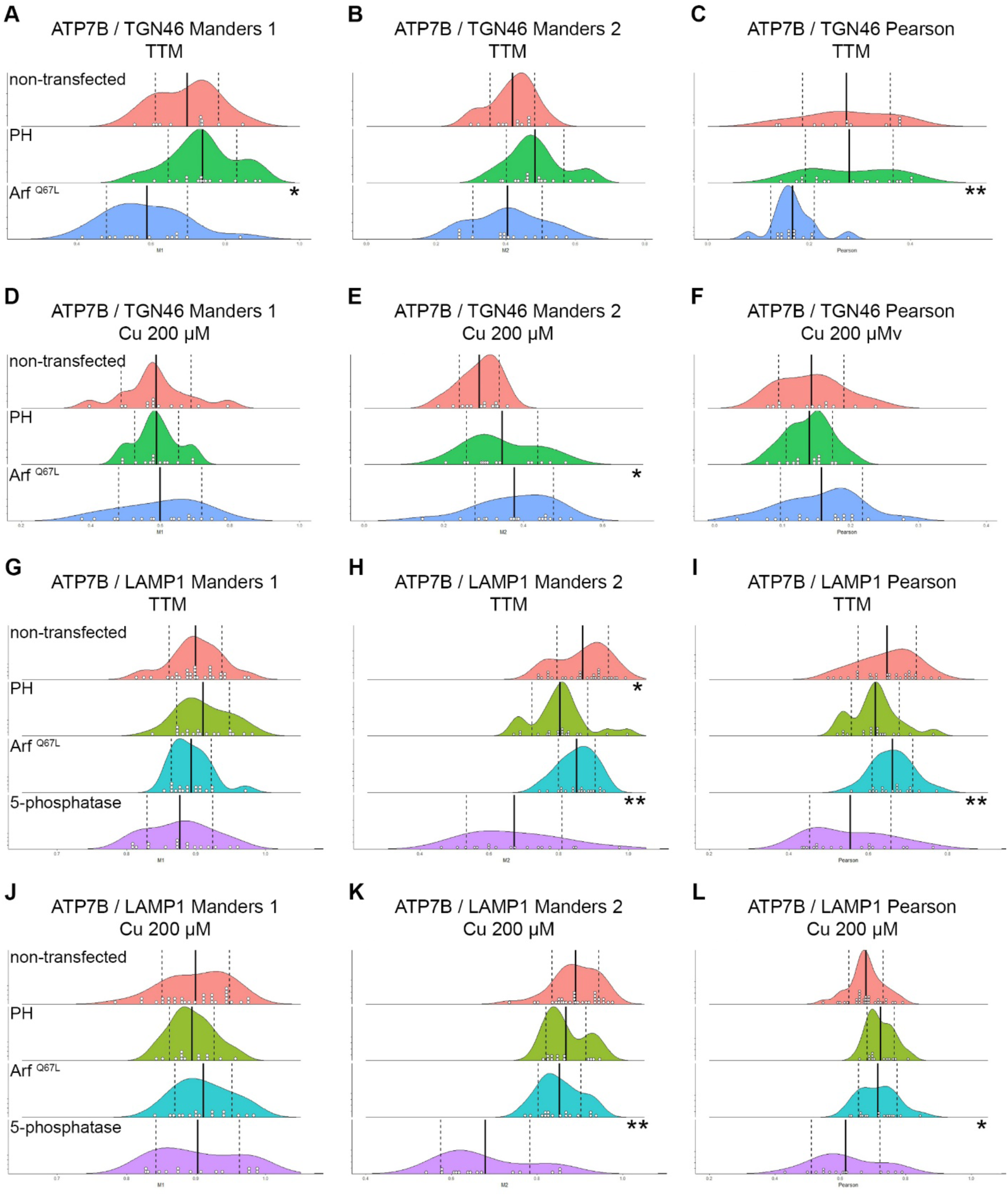
Colocalization of ATP7B with LAMP1 or TGN46 in response to PtdIns(4,5)2 modulation. HepG2 cells were transfected with Pleckstrin homology domain (PH) or Arf^Q67L^ then treated with TTM (low Cu) or 200μM Cu for 9 hours with cycloheximide added for the last hour. 3D colocalization analysis was used to produce a Menders and Pearson correlation coefficient for individual cells. Treatment means (solid line) and SD (dashed line) are shown. Individual cell values are indicated as a white circle (binned at 0.005). “*” indicates a p-value < 0.05 “**” indicates a p-value < 0.005.

**Figure 9.**
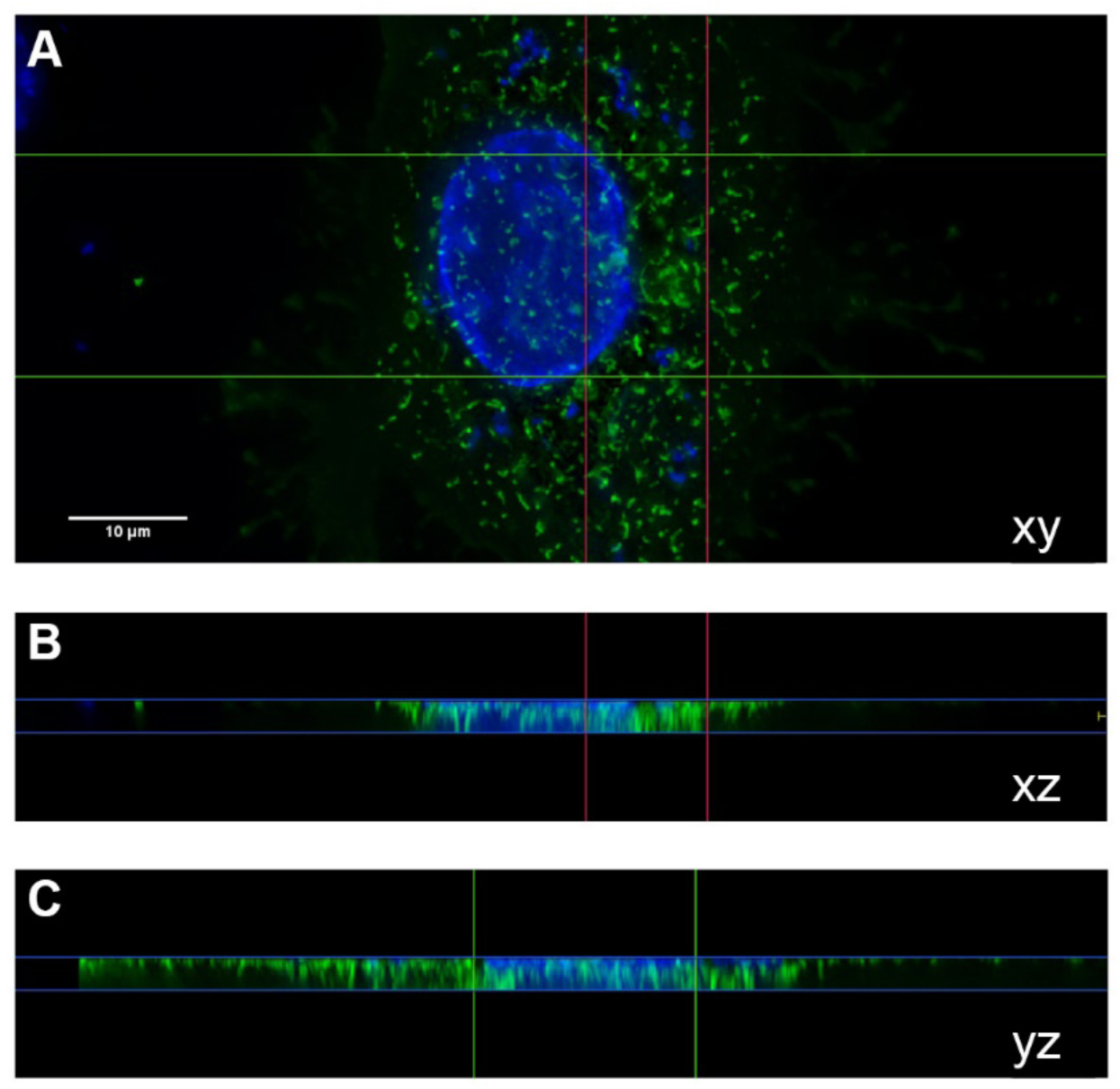
Intracellular localization of Pleckstrin homology domain in HepG2 cells. Cells were transfected with vectors containing GFP tagged Pleckstrin homology domain (PH-GFP) then imaged in 12-14 image stacks. Image stacks were deconvolved then visualized and analyzed using Huygens Professional.

## DISCUSSION

Prior studies found that knockdown of COMMD1 in various cell lines resulted in increased copper retention (Burstein et al., 2004; Miyayama et al., 2010; Spee et al., 2007). Furthermore, knockdown in in mouse Hepa1-6 cells prevented ATP7B retrograde trafficking from the periphery of the cell to the TGN when copper levels were reduced (Miyayama et al., 2010). Previous studies have also identified COMMD1 as a component of the CCC complex and suggested it functions at the recycling subdomain of the early endosome (Bartuzi et al., 2016; Phillips-Krawczak et al., 2015). The current model for retromer-independent cargo retrieval suggests that the WASH complex is first recruited to the EE, which then recruits the CCC complex through interactions with the FAM21 tail. From here, the CCC complex can then recruit the “retromer-like” complex retriever, which then interacts with SNX17 and its bound cargo, leading to retrieval of the cargo from degradation (see McNally and Cullen, 2018 for an in-depth review (McNally and Cullen, 2018)). Moreover, COMMD1’s specific function in ATP7B trafficking and thus in maintaining cellular copper homeostasis was until now elusive. However, the broader impacts of COMMD1-mediated trafficking for targets other than ATP7B are not fully understood, as COMMD1 appears to function as a pleiotropic cargo-directing adaptor.

Our current study determined that COMMD1 attenuation in HepG2 cells reduces the amount of ATP7B at the lysosome and TGN when trafficking is induced by high copper conditions **(Figs. 2**,**3)**. Additionally, COMMD1 attenuation leads to ATP7B accumulation in the EE **(Fig. 4)** and an overall increase in ATP7B abundance **(Fig. 5)**. Taken together, COMMD1 functions to ensure fidelity of ATP7B recycling or trafficking to the lysosome. Since ATP7B is directed to the cytoplasmic surface of the lysosome (Polishchuk et al., 2014), consistent with our observations (**Fig. 10)**, this targeting is not for degradation but to function in copper sequestration.

**Figure 10.**
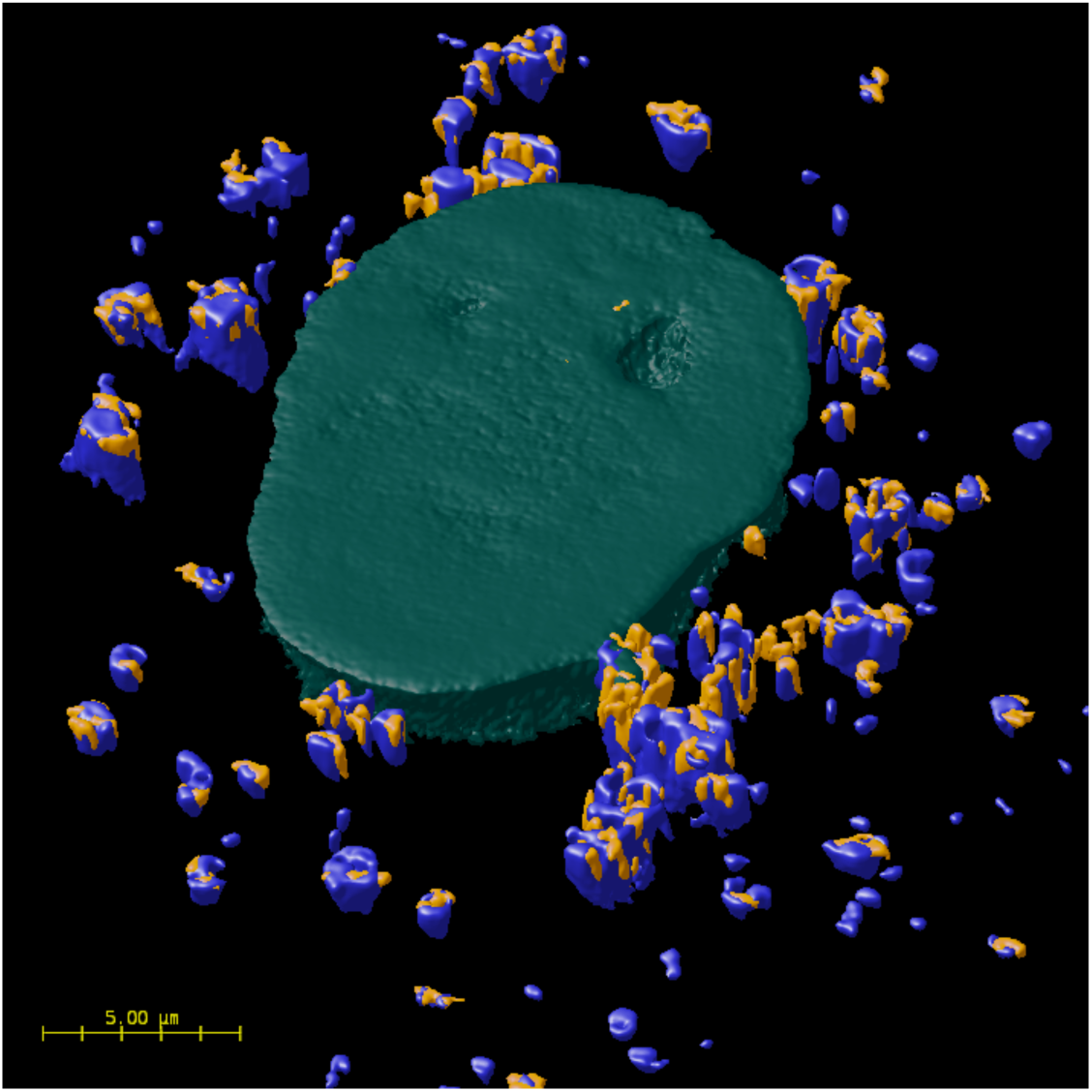
3D surface render of ATP7B (yellow), LAMP1 (blue), and the nucleus (cyan). ATP7B is observed localized at the surface of lysosomes. HepG2 cells were imaged in stacks of 12-14 image slices with z-steps of 0.198µm. Image stacks were deconvolved and a 3D surface render was created using Huygens Professional.

Determination of a specific role for COMMD1 in modulation of ATP7B trafficking provides new information about the cellular and biochemical functions of COMMD-containing trafficking complexes such as CCC and COMMANDER. A consistent pool of ATP7B colocalized with the TGN was observed, regardless of copper treatment. The portion of ATP7B colocalized with the lysosome increased when copper levels were increased **(Fig. 1)**. Under high copper conditions the cell must continue to produce cuproproteins while simultaneously managing and sequestering the excess copper load. It is likely the cell is upregulating ATP7B under these conditions and the additional ATP7B is directed to the lysosome. The data herein inform a model where ATP7B functions at the TGN for biosynthesis of cuproporteins and at the surface of the lysosome for sequestration of excess copper. In this model, ATP7B traffics between the TGN, the EE and the lysosome and its distribution is regulated at the EE by COMMD1 and its interaction with PtdIns(4,5)P_2_ **(Fig. 11)**.

**Figure 11.**
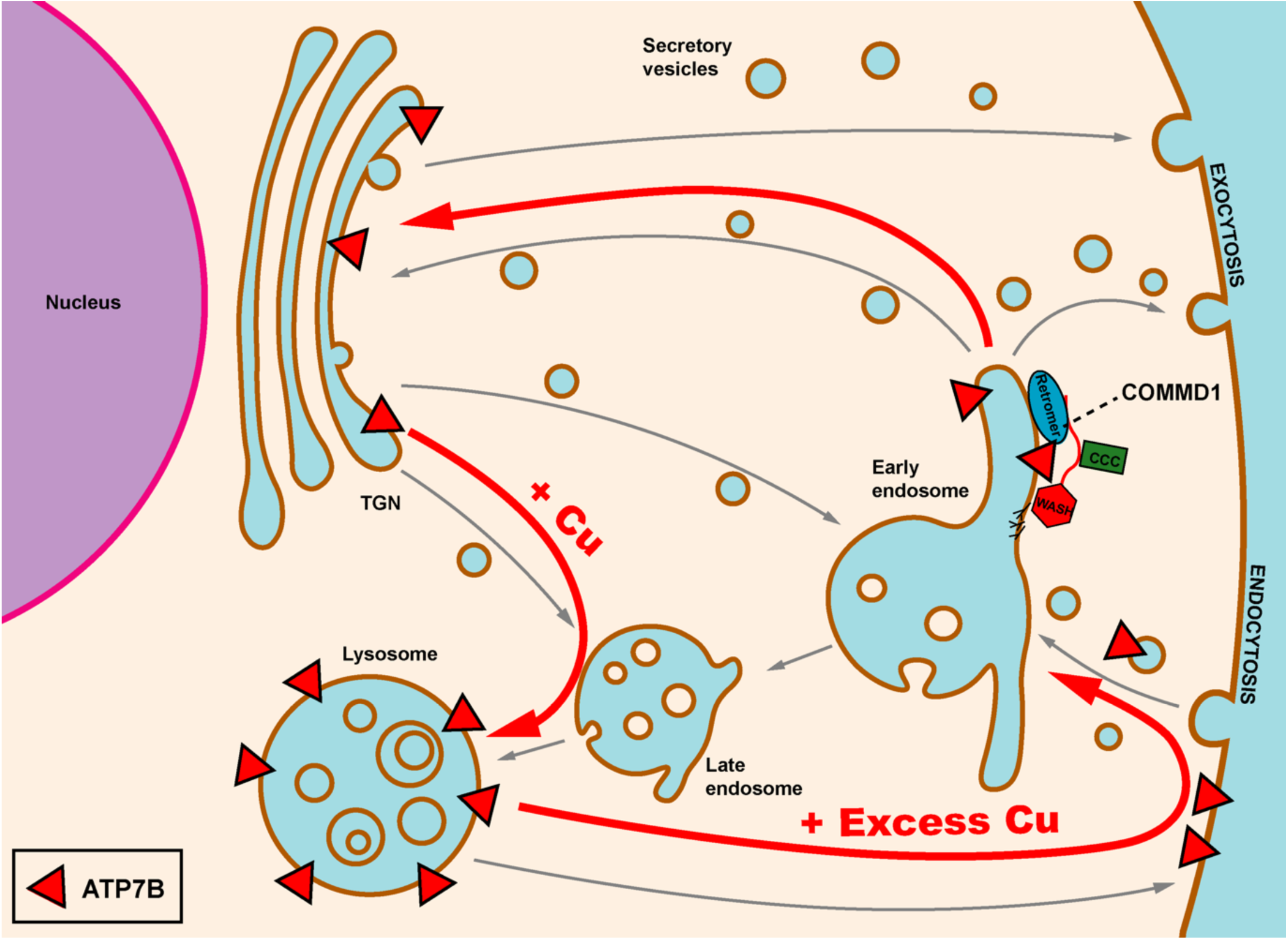
Schematic of ATP7B trafficking itinerary in hepatocytes. COMMD1 promotes ATP7B exit from the early endosome as part of the retromer/CCC complex.

Prior studies reported that COMMD1 interacts with phosphatidylinositols with a strong preference for PtdIns(4,5)P_2_ and to a lesser degree with PtdIns(4)P and PA (Burkhead et al., 2009). Other recent work suggested that COMMD1 binds non-specifically to negatively-charged phospholipids (Healy et al., 2018). However, our current experiments find that attenuation of COMMD1’s interaction with PtdIns(4,5)P_2_ results in altered ATP7B trafficking **(Fig. 6)**, supporting a distinct role for PtdIns(4,5)P_2_ in COMMD1-modulated trafficking. Specifically, overexpression of wildtype COMMD1 appears to direct ATP7B away from the lysosome. Whereas expression of the mutant variants, with decreased ability to bind PtdIns(4,5)P_2_, reduces the amount at the TGN. It may be that the interaction with PtdIns(4,5)P_2_ acts as the decision-making switch that directs ATP7B to the TGN or to function at the lysosome. Curiously, a current paradigm suggests that PtdIns(4,5)P_2_ is indicative of the PM and related processes (Balla, 2013). However, our observations with respect to COMMD1-mediated trafficking as well as the observation of PLCδ PH domain in association numerous intracellular vesicles suggests a broader cellular role. The impact of PLCδ PH on ATP7B trafficking is also suggestive of a specific role for this lipid in ATP7B trafficking, likely through COMMD1 action **(Fig. 8)**. Thus, in addition to the PM-related signaling, PtdIns(4,5)P_2_ may function as an important regulator at locations that include PM-derived components such as the EE. Taken together, this work begins to answer the question of how the cell maintains trafficking fidelity of ATP7B (through COMMD1 PtdIns(4,5)P_2_) and reveals a cellular and biochemical mechanistic model for COMMD function in cells. Though additional trafficking components have been identified in other studies, this work establishes a mechanism of action for COMMD1 and PtdIns in maintenance of cellular trafficking itineraries.

From a technical standpoint, this work emphasizes the importance of quantitative methods in studies of membrane protein trafficking. Here, an application is presented that illustrates the utility of quantitative immunofluorescence microscopy and statistical analysis. Additionally, these results demonstrate that our quantitative methods provide a refined view of subcellular protein distribution.

## MATERIALS AND METHODS

### Cell culture, plasmid transfection and copper treatment

Tet-on HepG2 cells (Clontech-Takara) were seeded on poly-lysine coated glass cover slips in 12-well plates at 0.7×10^5^ cells/well and grown in MEM/EBSS with 10% FBS and penicillin/streptomycin at 37°C with 5% CO_2_. Cells were then incubated for two days then transfected with specified plasmids or siRNA using TurboFect (ThermoFisher) followed by overnight incubation. Cells were examined for morphology consistent with HepG2 cells. Routine DAPI staining as part of immunofluorescence microscopy ensured that the cells were free of mycoplasma contamination.

Copper treatments: cells were treated with 10 µM CuCL_2_, 100 µM CuCL_2_ 200 µM CuCL_2_ (‘CU’ label in figures) or 10 µM tetrathiomolybdate (‘TTM’ label) for 9 hours. Cycloheximide was added for the last hour to prevent new protein synthesis. The plating, growth and transfection procedure allowed sufficient time for HepG2 cells to polarize(Sormunen et al., 1993). Experiments were repeated three or more times.

COMMD1 expression was attenuated using esiRNA (Sigma-Aldrich, St. Louis, Mo) at concentrations of 1nM, 5nM, 10nM, 15nM, and 20nM or with 10nM nontarget control RNA.

### Plasmids and antibodies

The initial cloning of the COMMD1 cDNA sequence is described in Burkhead et al., (2009). The COMMD1 sequence was amplified by polymerase chain reaction using the *Pfu* DNA polymerase (Roche, Basel, Switzerland), using an upstream primer to add a *Hind*III restriction site and retaining the *Sal*I downstream of the coding sequence. This insert was ligated into the *Hind*III and *Sal*I sites in the peGFP-C1 plasmid (Clontech). The eGFP and COMMD1 coding sequence was subcloned to pTre-TIGHT-Bi (Clontech) by introducing a *Bam*H1 site upstream of the eGFP start codon and ligating the eGFP-COMMD1 insert into the *Bam*H1 and SalI sites of pTRE-Tight-BI (Clontech). The insert sequence was verified by dideoxy-Sanger sequencing. T174M and K167E/K173E mutations were introduced to the same construct by Quickchange site-directed mutagenesis (Agilent) and verified by dideoxy-Sanger sequencing. Overexpression of COMMD1 and variants used the pTre-TIGHT-Bi plasmids containing wild type, T174M or K167E/K173E COMMD1 transfected into HepG2 cells as described above. GFP-COMMD1 expression was induced by inclusion of 750 nM doxycycline in growth media. For *E. coli* expression, COMMD1-T174M and COMMD1-K167/173E were synthesized and inserted into pET32 with an N-terminal His6 tag and thrombin cleavage site as in previous work with pET28b (Burkhead et al., 2009).

The following plasmids were generously donated: HA-AfrQ67L was provided by J. Donaldson HA-AfrQ67L (Donaldson, 2003) (NHLBI/NIH, Bethesda, MD), MYC-5-ptase IV was provided by P. Majerus MYC-5-ptase IV (Kisseleva et al., 2002) (Washington University School of Medicine, St. Louis, Mo), PH-GFP was provided by T. Balla (Várnai and Balla, 1998) (NIH, Bethesda, MD).

Primary antibodies and concentrations used for immunofluorescence microscopy were: 1:750 Rabbit anti-ATP7B (Abcam, Cambridge, MA), 1:1000 Mouse anti-LAMP1 (Developmental Studies Hybridoma Bank, Iowa City, Iowa), 1:300 Sheep anti-TGN46 (Acris Antibodies, Inc., San Diego, CA), 1:300 Mouse anti-Golgin-97 (Invitrogen, Carlsbad, CA), 1:500 Goat anti-VPS35 (Abcam, Cambridge, MA), 1:500 Rat anti-HA (Roche Diagnostics, Indianapolis, IN), 1:500 Rat anti-C-Myc (BioRad, Hercules, CA), Mouse anti-COMMD1 (Novus Biologicals, Littleton, CA). Secondary antibodies were conjugated to Alexa 488 and 647 (Invitrogen, Carlsbad, CA) and Dylight 550 (Novus Biologicals, Littleton, CO).

### Immunofluorescence staining

Cells grown on lysine-coated circle coverslips were fixed in cold 4% paraformaldehyde in PBS for 20 minutes in the dark, followed by blocking and permeabilization in BP buffer (3% BSA, 50mM NH_4_Cl, 0.1% Saponin in PBS with sodium azide) at 4°C overnight. Cells were then incubated with primary antibodies diluted in BP buffer for two hours at room temperature in a dark humidified chamber. After three 10-minute washes in BP buffer, the cells were incubated in secondary antibodies diluted in BP buffer for 45 minutes at room temperature in a dark humidified chamber. This was followed by three washes in BP buffer, a 10-minute incubation in DAPI diluted 1:1000 in PBS, and two final washes in PBS. Cover slips were then rinsed in water and mounted to slides with Fluoromount-G (Southern Biotech).

### Immunofluorescence microscopy and 3D deconvolution

Cells were imaged using a Leica DM6000 widefield microscope with a 63x 1.40 − 0.60 oil objective and Photometrics CoolSNAP MYO CCD camera. Cells with intact nuclei were selected for imaging. For cells expressing GFP constructs, GFP expression was assessed and only cells with moderate expression and intact nuclei were imaged. Images were acquired as four channel 16-bit 1940×1460 (0.072 × 0.072 µm pixels) stacks of 12-14 image slices with z-steps of 0.198µm. Image stacks were then deconvolved using LAS-X 3D deconvolution using 10 iterations, an automatically generated point spread function, background removal, and intensity rescaling to compensate for any bleaching that occurred during image acquisition of the stack.

### Colocalization and statistical analysis

The deconvolved image stacks were analyzed with Huygens Professional (Scientific Volume Imaging) for colocalization analysis. ROI masks were created for each image to select cells of interest. Background for the two channels of interest was estimated using Huygens Optimized method, which is a variation of Costes method but uses the entire histogram rather than the linear regression line to iteratively determine the background level. Pearson and Manders M1 and M2 correlation coefficients were then generated for each ROI. (For an in-depth Review of Pearson’s correlation and Manders overlap coefficients see *A practical guide to evaluating colocalization in biological microscopy* (*2011*) (Dunn et al., 2011). In brief, Pearson’s correlation coefficient is a relatively simple statistic that measures the pixel-by-pixel covariance in the signal intensity of two images. The formula is for an image consisting of both a red and a green channel where *R*_*i*_ and *G*_*i*_ refer to the intensity values of the channels of *pixel i*, and 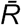 and 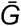 refer to the mean intensities of the channels across the entire image.

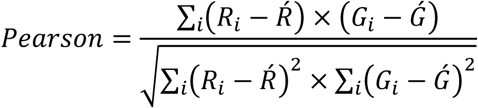

Pearson’s correlation is used to measure the degree to which a linear relationship can be used to explain the variability in pixel intensity in the red and green channels. Thus, Pearson is a useful measure of association in a biological system when the relationship between the two probes is linear (Dunn et al., 2011). Here we use Pearson to compare the degree to which probe association changes from one treatment to the next.

The Manders coefficient is a more direct and intuitive measure of the fraction of one protein that colocalizes with a second protein. Manders generates two coefficients for an image consisting of both a red and a green channel; M1 is the fraction of pixels in R that colocalize with pixels in G independent of intensity, and M2 is the fraction of pixels in G that colocalize with pixels in R independent of intensity. These values are calculated as:

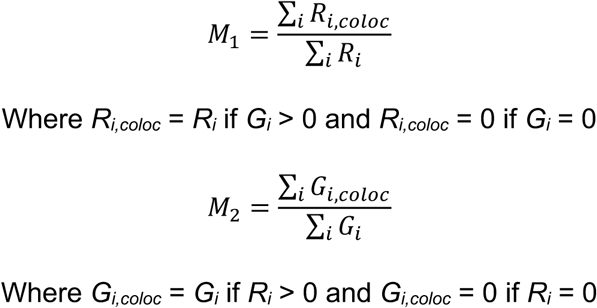

It is important to note that Manders values are calculated independently of the probes’ intensities and therefore strictly measures co-occurrence(Dunn et al., 2011). For our purposes, the ‘green’ channel (G) will represent ATP7B and the ‘red’ (R) channel will represent the probe we are measuring against (TGN, lysosome or VPS35).

We used JMP Pro 13 (SAS Institute) for statistical analysis. Power analysis based on preliminary experiments was used to determine an appropriate sample size. For each treatment, 15 to 20 cells were analyzed (biological replicates). Data was compiled and cells in a given treatment were treated as a population. Student’s T-test was used for direct comparisons between two treatments. Tukey’s HSD test was used for multiple comparisons to test for differences between all tested pairs. Dunnett’s test was used to compare multiple treatments individually against a single control. Density plots were created in *RStudio* using the plyr (Wickham, 2011) and ggplot (Wickham and Chang, 2016) packages to better visualize the distribution of cell populations.

### Microsomal membrane preparation, Western blot and densitometry

Cells were grown and treated as above. Cells were washed with cold PBS then placed at −80°C for 10 min. The cells were then resuspended in 1.0 mL homogenization buffer (25mM Tris-HCl pH 7.4, 250 mM sucrose, 1 mM PMSF and protease inhibitor cocktail (Sigma)) and homogenized with a Dounce homogenizer (40 strokes with a tight pestle). Lysed cells were then centrifuged at 700×*g* for 10 minutes at 4°C to pellet cell debris and nuclei, 3,000×*g* for 10 minutes at 4°C to pellet mitochondria, and finally at 20,000×*g* for 30 minutes at 4°C to pellet microsomal membranes. The supernatant was removed and concentrated using a 0.5 mL Pierce Concentrator with a 10K MWCO (Prod# 88513, ThermoFisher) and used for COMMD1 analysis. The pellet was resuspended in 40 µL NP40 buffer (50 mM Tris pH 8.0, 150 mM sodium chloride, 0.5% NP-40, 1 mM PMSF, and protease inhibitor cocktail) and used for ATP7B analysis. Sample protein concentrations were determined via Pierce BCA Protein Assay (Prod# 23227) and used to determine equal loading. ATP7B and COMMD1 abundance was assayed by sodium dodecyl sulfate-polyacrylamide gel electrophoresis (SDS-PAGE) analysis using 15% and 7.5% Acrylamide gels for COMMD1 and ATP7B respectively. Proteins were then transferred to a PVDF membrane and visualized with monoclonal antibodies against ATP7B and COMMD1. After imaging, blot was stained with Coomassie and imaged. Protein abundance was determined by densitometry using ImageJ and the Coomassie blot was used to correct for unequal loading.

### *In-vitro* PtdIns(4,5)P_**2**_ binding to COMMD1

Recombinant COMMD1 and variants T174M or K167E/K173E) were expressed and purified essentially as described in Burkhead et al., (2009). The His6 tag was cleaved by addition of GST-thrombin to the refolded protein. Thrombin was removed by incubation with glutathione-sepharose beads and the cleaved tag was removed by dialysis. Purified proteins were dialyzed to 20mM HEPES, 50mM Potassium Acetate, 1.0mM EDTA, and 0.05% CHAPS.

An intrinsic fluorescence quenching binding assay was prepared using PtdIns(4,5)P_2_ (“PIP2”, 1,2-dihexanoyl sodium salt) (Cayman Chemical) as a ligand. Protein concentration was determined using OD_280_ with adjusted extinction coefficients (OD280 of the native protein vs. OD280 of denatured (6 M Gdn-HCl). The initial point of the assay contained 50 nM of COMMD1 or mutant and with final volume brought to 2mL. PIP2 was then titrated in 1µL to 10µL increments (total volume increase at end of assay was 35µL) and allowed 5 minutes equilibration time for each addition. Concentrations of PIP2 were 0µM, 50µM, 100 µM, 150 µM, 200 µM, 250 µM, 300 µM, 400 µM, 500 µM, 750 µM, 1000 µM, 3500 µM, and 8500 µM. Intrinsic fluorescence was measured with a Horiba Fluoromax-3 spectrometer using an excitation of 280 nm and emission scans from 300-400 nM. Three scans were averaged for each data point and the assay was performed in triplicate for each protein variant. Peak at emission at 332 nm was used for calculations (*n*=3 for each variant). Graphs were generated in GraphPad Prism 7 to fit with nonlinear curve with equation Binding-Saturation and One site-Total.

## Acknowledgements

This work was supported by National Science Foundation grant MCB-1411890 to JLB and an Institutional Development Award (IDeA) from the National Institute of General Medical Sciences of the National Institutes of Health under grant number P20GM103395. We thank Ezra Burstein for helpful comments and insight, as well as Elena Buglo, Caitlin Kollander, Nicholas Braman and Jessica Schwartz for lab assistance in development of the project.

